# Novel repolarisation metric predicts arrhythmia origin and clinical events in ARVC and Brugada Syndrome

**DOI:** 10.1101/123422

**Authors:** CA Martin, M Orini, N Srinivasan, J Bhar-Amato, S Honarbakhsh, A Chow, MD Lowe, RD Simon, PM Elliott, P Taggart, PD Lambiase

**Affiliations:** Barts Heart Centre, Barts Health NHS Trust, West Smithfield, London EC1A 7BE, UK; Institute of Cardiovascular Science, University College London, Gower Street, London WC1E 6BT, UK

**Author notes:** Equal contributors (CA Martin, M Orini). Address for correspondence: Professor Pier D. Lambiase, Cardiology Dept, Barts Heart Centre, Barts Health NHS Trust, West Smithfield, London, EC1A 7BE, UK. Email addresses.

**Keywords:** Repolarization, Arrhythmogenic Right Ventricular Cardiomyopathy, Brugada Syndrome, ventricular tachycardia, ablation, risk stratification

## Abstract

**Background:** Initiation of re-entrant ventricular tachycardia (VT) involves complex interactions between activation (AT) and repolarization times (RT). The re-entry vulnerability index (RVI) is a recently proposed activation-repolarization metric designed to quantify tissue susceptibility to re-entry.

**Objectives:** The study aimed to test the feasibility of an RVI-based algorithm to predict the exit site of VT and occurrence of clinical events.

**Methods:** Patients with Arrhythmogenic Right Ventricular Cardiomyopathy (ARVC) (n=11), Brugada Syndrome (BrS) (n=13) and focal RV outflow tract VT (n=9) underwent programmed stimulation with unipolar electrograms recorded from a non-contact array. The distance between region of lowest RVI and site of VT breakout (D_min_), and global minimum RVI (RVIG) were computed to assess prediction of site of VT breakout and occurrence of clinical events, respectively.

**Results:** Lowest values of RVI, representing sites of highest susceptibility to re-entry, co-localised with site of VT breakout in ARVC/BrS but not in focal VT and Dmin values were lower in ARVC/BrS. ARVC/BrS patients with inducible VT had lower RVIG than those who were non-inducible or those with focal VT. Patients were followed up for 112 ± 19 months; those with clinical VT events had lower RVIg than those without VT or those with focal VT.

**Conclusions:** The proposed methodology based on RVI localises the origin of re-entrant but not focal ventricular arrhythmias and predicts clinical events. This index could be applied to target ablation for arrhythmias which are difficult to induce or are haemodynamically unstable and also risk stratify patients for ICD prophylaxis.

Abbreviations list

AT: Activation time
RT: Repolarization time
RVI: Re-entry vulnerability index
ARVC: Arrhythmogenic Right Ventricular Cardiomyopathy
BrS: Brugada Syndrome
VT: Ventricular tachycardia
RVOT: Right ventricular outflow tract
EPS: Electrophysiological study
VERP: Ventricular effective refractory period

## Introduction

A significant challenge remains in the prediction of ventricular tachycardia (VT) circuits, especially in disorders of diffuse fibrosis or where the functional characteristics of the substrate play a critical role in the initiation of re-entry. This is reflected in Arrhythmogenic Right Ventricular Cardiomyopathy (ARVC) and Brugada Syndrome (BrS), where fibrosis in the right ventricle can be especially difficult to image due to the diffuse epicardial nature of the disease and the limitations of magnetic resonance imaging (MRI) resolution in the thin RV free wall (1–3). Both the prediction of ventricular arrhythmic events and targeting of ablation remain a demanding problem in these conditions due to the diffuse substrate and the presence of multiple VT morphologies or ventricular fibrillation (VF) leading to haemodynamic compromise.

Recently a novel spatial metric has been developed termed the Re-entry Vulnerability Index (RVI), used to locate regions of tissue susceptible to re-entry (4, 5) (Figure 1). The RVI is based on previous experimental studies which quantitatively assessed the likelihood of excitation wavefront-waveback interactions between two points spanning a line of conduction block following a premature stimulus (6). The algorithm has subsequently been shown to accurately identify the region of a macro re-entrant circuit in two animal models of ischaemia and a single clinical case of ischaemic cardiomyopathy (5). Pilot simulations of RVI maps within a rabbit ventricular model (7) suggested that the metric was able to reliably identify exit sites associated with re-entrant circuits for different scar morphologies.

**Figure 1.**
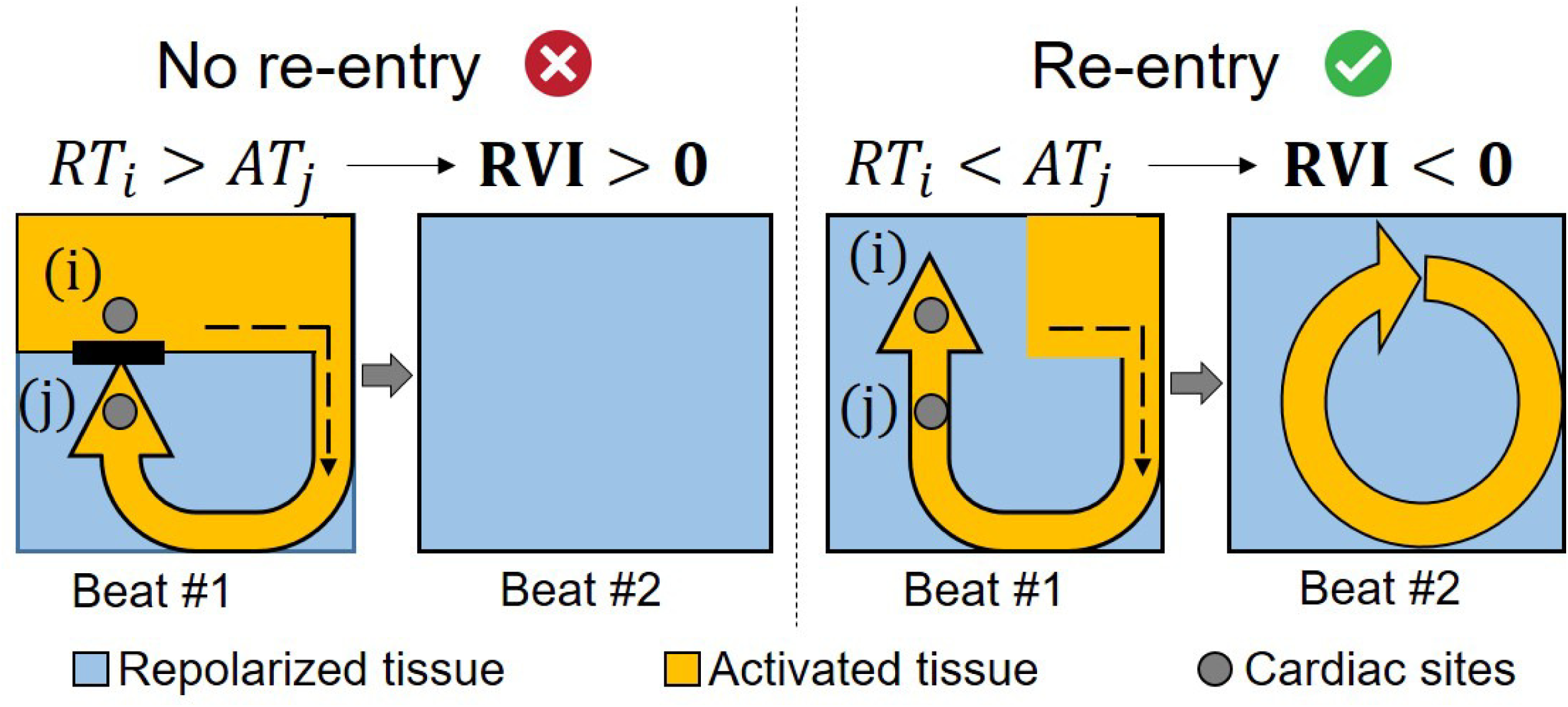
The time interval between the arrival of the wave at the exit site and the regaining of excitability (repolarisation) of the healthy tissue just proximal to it is a critical factor in determining the success of such a circuit to block or sustain re-entry.

The aim of this study was to test the feasibility of RVI mapping in the RV for patients with BrS and ARVC. We hypothesised that (i) regions of low RVI may predict sites vulnerable to re-entry and therefore VT initiation and (ii) the global RVI may predict the propensity to clinically important ventricular arrhythmias to aid risk stratification in the prevention of sudden cardiac death.

## Methods

### Ethics statement

The research was approved by University College London Hospitals Ethics Committee A (08/H0714/97). Prior written informed consent to participate in this study was obtained from all participants.

### Patient selection

38 patients aged 18–65 years were prospectively recruited to participate and informed consent was obtained. Inclusion criteria were a diagnosis of ARVC by modified task force criteria (including familial or genetic criteria) or a diagnosis of BrS based on type I ECG pattern (either spontaneous or provoked) with one of the following: documented VF, polymorphic VT, syncope or a family history of SCD. Patients with benign RV outflow tract (RVOT) ectopy or VT were used as a control cohort. All patients underwent detailed assessment with resting ECGs, signal averaged ECGs (SAECG) and echocardiography/MRI. Patients with a clinical suspicion of BrS underwent ajmaline provocation testing.

Following electrophysiological study (EPS), all patients had at least 50 months of follow up (mean = 112 ± 19 months). Patients were seen annually with clinical assessment, resting ECG, 24 hour tape or device interrogation and serial imaging if appropriate. A clinical arrhythmic event was defined as VF or haemodynamically unstable VT either terminating spontaneously or by anti-tachycardia pacing (ATP) /ICD shock.

The RVOT VT group had normal resting and SAECGs, only unifocal ectopy or VT, structurally normal hearts on echocardiogram/MRI and no family history of SCD. They were therefore classified as RV outflow tract focal VT. In order to minimize the likelihood of a patient with their first presentation of hitherto clinically silent cardiomyopathy or channelopathy being misassigned a diagnosis of benign RVOT ectopy, focal VT patients were excluded if they had a recurrence of ventricular ectopy/tachycardia post ablation.

### Genetic testing

Whole blood samples were obtained from participating BrS and ARVC subjects. Genomic DNA was extracted using a commercially available DNA extraction kit (QIAamp DNA Blood mini kit, Qiagen). ARVC cases were screened for mutations in desmoplakin, plakoglobin, plakophilin-2, desmoglein-2 and desmocollin-2, and BrS cases for mutations in SCN5A.

### Electrophysiological mapping

The procedure for non-contact mapping has been previously described (8, 9). The 64 electrode non-contact array (St Jude Medical, USA) was placed in the RVOT via the left or right femoral vein under conscious sedation and was positioned within 4cm of the endocardial surface to obtain accurate non-contact unipolar electrogram data. The geometry of the RV endocardium was created with a steerable quadrapolar mapping or ablation catheter.

Programmed stimulation was performed from the RV apex. 3 minutes of steady state pacing at 600 ms coupling intervals was followed by an electrical stimulation protocol, consisting of an 8 beat train of pulses (S1) at 600 ms coupling interval followed by a single, premature stimulus (S2). The S_1_S_2_ coupling interval was reduced sequentially from 400 ms by 10 ms until ventricular refractoriness (VERP). A VT stimulation protocol followed, consisting of an 8 beat drivetrain from the RV apex at 600 and 400 ms baseline cycle length followed by S_2_ until VERP or 200 ms coupling interval was reached. This was repeated for each baseline drivetrain adding an S_3_ & S_4_.

### Offline analyses

The non-contact mapping data consisting of 256 unipolar electrograms were collected with a recording bandwidth of 0-300 Hz in all cases and measurements made with a filter band width of 0.1-25 Hz. Electrograms were exported and analysed using semi-automated custom software running in Matlab (The Mathworks Inc., MA, USA) (10, 11). During pacing, the time from pacing artefact to the steepest negative deflection (dV/dt_min_) was used as the local activation time (AT). The classical (Wyatt) method was used to calculate the repolarisation time (RT) (Figure 2) (12).

**Figure 2.**
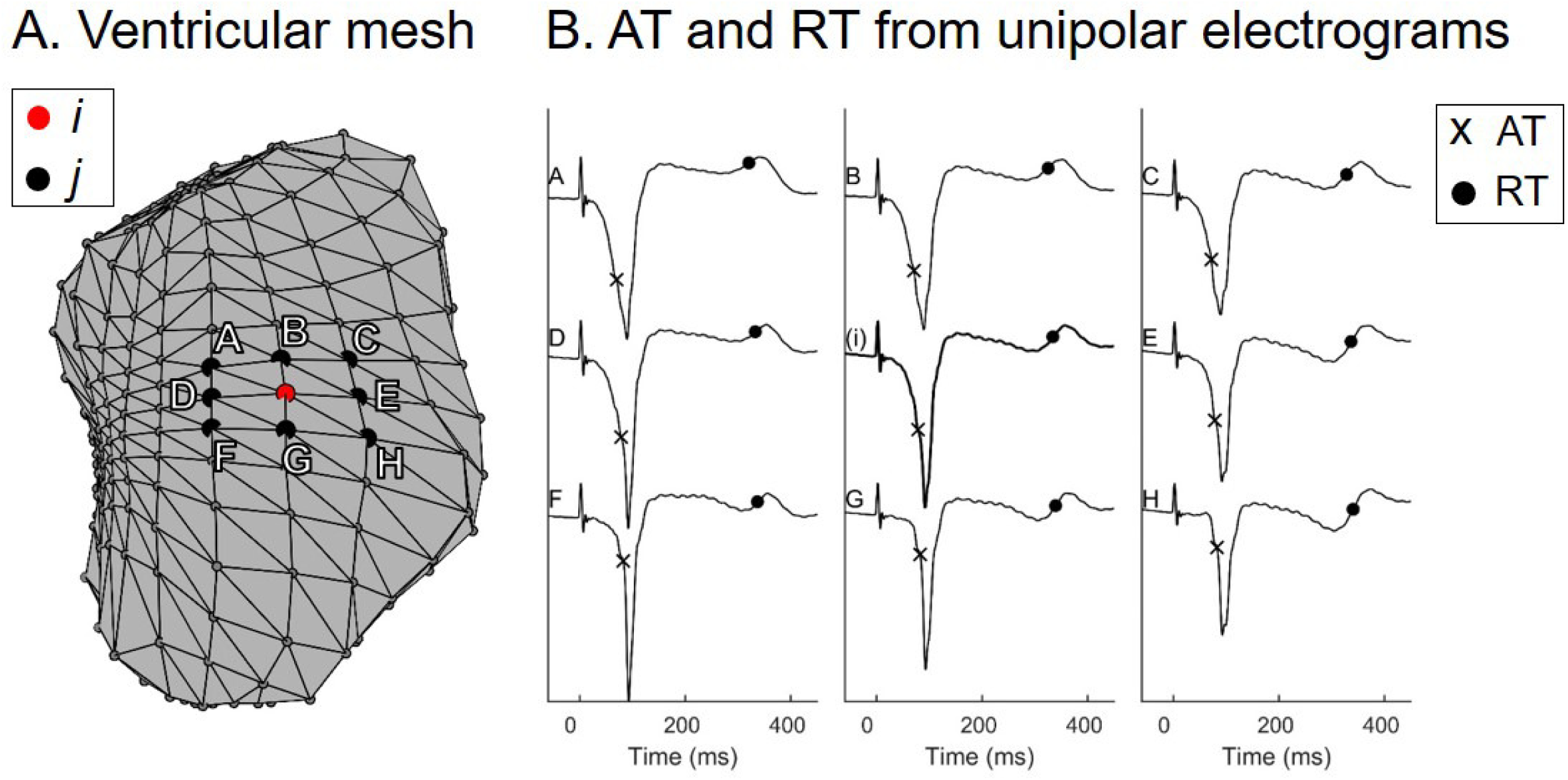
Calculation of RVI: Neighbouring sites to an electrogram recording are identified within a pre-defined radius. A: Representative ventricular mesh from the clinical system, showing a site *i* and its closest neighbours *j*. B: RVI at site *i* is the minimum difference between RT at site *i* ATs at sites *j*. AT and RT are represented as crosses and circles in panel B.

The RVI of each recording location *i*, RVI_i_, was calculated taking the minimum difference between RT at site *i*, RT_i_, and AT at neighbouring sites *j* comprised within a 20mm radius, AT_j_,: 
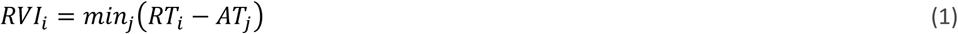

Since vulnerability increases for lower RVI values, the definition of RVI_i_ as the minimum difference between RT_i_ and AT_j_ ensured high spatial sensitivity to activation/repolarization abnormalities that may promote re-entry. The patient-specific location of the 256 ventricular sites from the clinical mapping system was then used to create an RVI colour map.

A global index of re-entry vulnerability, RVI_G_, was derived from the map by subtracting the median RVI from the 10^th^ percentile of the RVI distribution, P_10%_(RVI_i_):

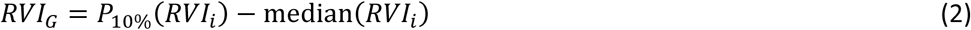

This index is a robust version of the minimum RVI corrected to reduce cycle length dependency and it quantifies the global vulnerability of each patient.

For patients who developed VT during the VT stimulation protocol, the breakout site was identified as the earliest site of activation of the first VT beat (VT_AT_). The distance (*D_min_*) between the region of lowest RVI (including the bottom 5% of all RVI values) and the region of earliest activation during VT (including sites that activate within 5 and 20 mm from the site of earliest activation) was computed for each patient.

In order to provide consistent RVI measurements between hearts, for RVI_G_ calculations the unipolar electrograms of a single S2 beat with approximately the same S_1_S_2_ coupling interval (348 ± 17 ms) for each patient were used. For *D_min_* calculations where region of lowest RVI within each heart was of primary interest, the S_2_ or where applicable S_3_ beat immediately preceding VT initiation was used.

## Results

### 1. Patient demographics

Demographics of all study subjects (13 BrS (54% male, age 53 ± 13 years), 11 ARVC (64% male, age 62 ± 12 years) and 9 focal RVOT VT (33% male, age 51 ± 8 years)) are shown in Table 1. In the BrS patients, 2 had a VF arrest at presentation and 1 had unexplained syncope; 3 were diagnosed with a BrS type 1 ECG incidentally and the remainder were diagnosed through family screening. Six (46%) had a positive VT simulation study with induced sustained VT/VF. None of the patients who were non-inducible at EPS had a clinical event. 5 BrS patients (38%) had an ICD; 2 were implanted for secondary prevention in the patients with VF arrests and 3 were implanted for primary prevention following inducible VT at EPS and other high risk features (1 further patient with inducible VT declined ICD).

**Table 1.**
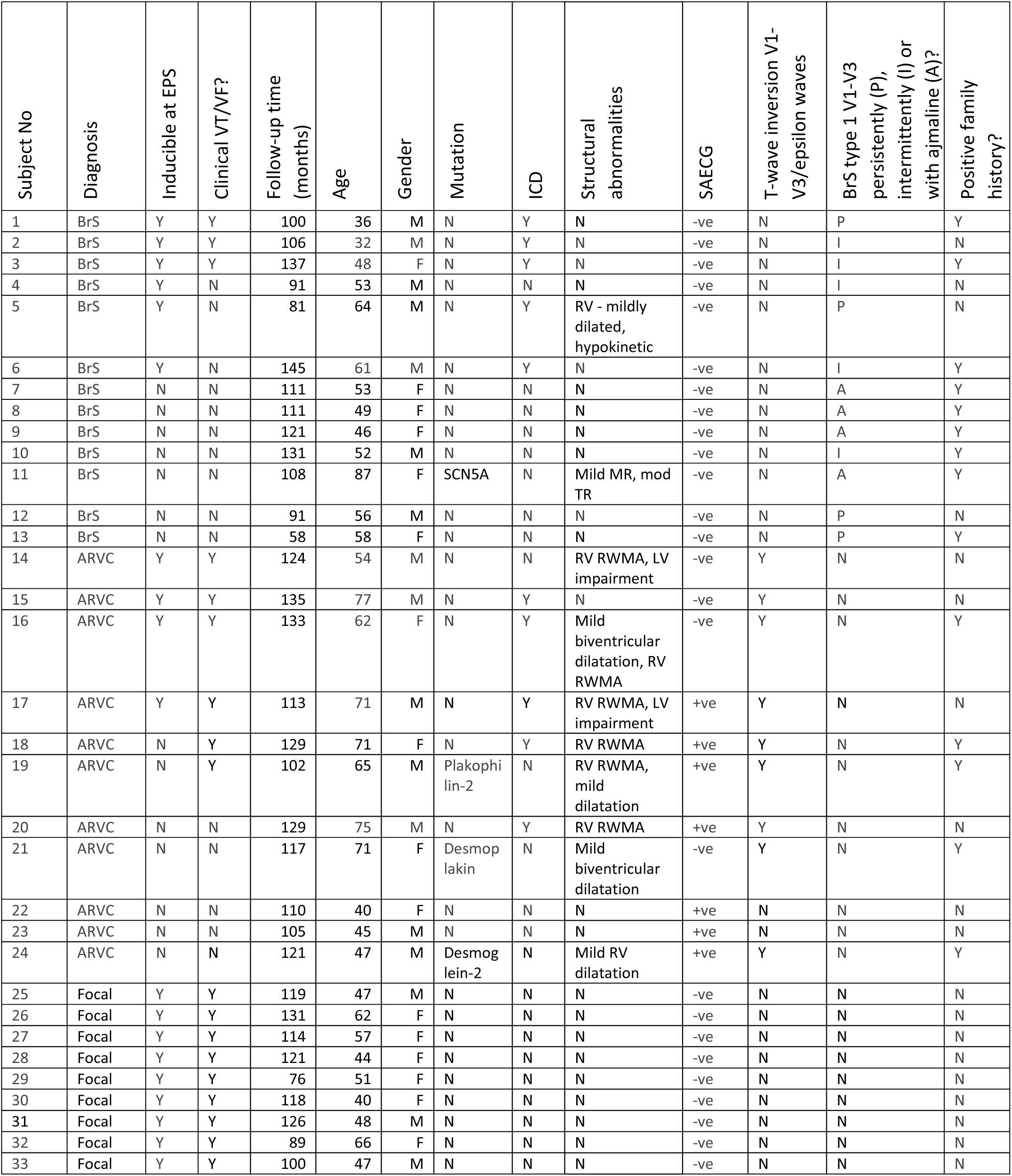
Table showing demographics of study participants. RWMA = regional wall motion abnormalities, MR = mitral regurgitation, TR = tricuspid regurgitation, LV = left ventricle.

| Subject No | Diagnosis | Inducible at EPS | Clinical VT/VF? | Follow-up time (months) | Age | Gender | Mutation | ICD | Structural abnormalities | SAECG | T-wave inversion V1-V3/epsilon waves | BrS type 1 V1-V3 persistently (P), intermittently (I) or with ajmaline (A)? | Positive family history? |
| --- | --- | --- | --- | --- | --- | --- | --- | --- | --- | --- | --- | --- | --- |
| 1 | BrS | Y | Y | 100 | 36 | M | N | Y | N | -ve | N | P | Y |
| 2 | BrS | Y | Y | 106 | 32 | M | N | Y | N | -ve | N | I | N |
| 3 | BrS | Y | Y | 137 | 48 | F | N | Y | N | -ve | N | I | Y |
| 4 | BrS | Y | N | 91 | 53 | M | N | N | N | -ve | N | I | N |
| 5 | BrS | Y | N | 81 | 64 | M | N | Y | RV - mildly dilated, hypokinetic | -ve | N | P | N |
| 6 | BrS | Y | N | 145 | 61 | M | N | Y | N | -ve | N | I | Y |
| 7 | BrS | N | N | 111 | 53 | F | N | N | N | -ve | N | A | Y |
| 8 | BrS | N | N | 111 | 49 | F | N | N | N | -ve | N | A | Y |
| 9 | BrS | N | N | 121 | 46 | F | N | N | N | -ve | N | A | Y |
| 10 | BrS | N | N | 131 | 52 | M | N | N | N | -ve | N | I | Y |
| 11 | BrS | N | N | 108 | 87 | F | SCN5A | N | Mild MR, mod TR | -ve | N | A | Y |
| 12 | BrS | N | N | 91 | 56 | M | N | N | N | -ve | N | P | N |
| 13 | BrS | N | N | 58 | 58 | F | N | N | N | -ve | N | P | Y |
| 14 | ARVC | Y | Y | 124 | 54 | M | N | N | RV RWMA, LV impairment | -ve | Y | N | N |
| 15 | ARVC | Y | Y | 135 | 77 | M | N | Y | N | -ve | Y | N | N |
| 16 | ARVC | Y | Y | 133 | 62 | F | N | Y | Mild biventricular dilatation, RV RWMA | -ve | Y | N | Y |
| 17 | ARVC | Y | Y | 113 | 71 | M | N | Y | RV RWMA, LV impairment | +ve | Y | N | N |
| 18 | ARVC | N | Y | 129 | 71 | F | N | Y | RV RWMA | +ve | Y | N | Y |
| 19 | ARVC | N | Y | 102 | 65 | M | Plakophilin-2 | N | RV RWMA, mild dilatation | +ve | Y | N | Y |
| 20 | ARVC | N | N | 129 | 75 | M | N | Y | RV RWMA | +ve | Y | N | N |
| 21 | ARVC | N | N | 117 | 71 | F | Desmoplakin | N | Mild biventricular dilatation | -ve | Y | N | Y |
| 22 | ARVC | N | N | 110 | 40 | F | N | N | N | +ve | N | N | N |
| 23 | ARVC | N | N | 105 | 45 | M | N | N | N | +ve | N | N | N |
| 24 | ARVC | N | N | 121 | 47 | M | Desmoglein-2 | N | Mild RV dilatation | +ve | Y | N | Y |
| 25 | Focal | Y | Y | 119 | 47 | M | N | N | N | -ve | N | N | N |
| 26 | Focal | Y | Y | 131 | 62 | F | N | N | N | -ve | N | N | N |
| 27 | Focal | Y | Y | 114 | 57 | F | N | N | N | -ve | N | N | N |
| 28 | Focal | Y | Y | 121 | 44 | F | N | N | N | -ve | N | N | N |
| 29 | Focal | Y | Y | 76 | 51 | F | N | N | N | -ve | N | N | N |
| 30 | Focal | Y | Y | 118 | 40 | F | N | N | N | -ve | N | N | N |
| 31 | Focal | Y | Y | 126 | 48 | M | N | N | N | -ve | N | N | N |
| 32 | Focal | Y | Y | 89 | 66 | F | N | N | N | -ve | N | N | N |
| 33 | Focal | Y | Y | 100 | 47 | M | N | N | N | -ve | N | N | N |

In the ARVC patients, 1 had episodes of sustained VT at presentation, 7 were diagnosed after investigation for palpitations and 3 were diagnosed through family screening. As well as the patient with clinical VT at presentation, another 5 patients had episodes of symptomatic VT during follow-up that was either sustained in patients without a device (n=2), or treated by ICD by ATP (n=3). Four patients (36%) had a positive VT stimulation study; all 4, plus a further 2 who had been negative at EPS, were the patients with clinical events. Five (45%) ARVC patients had an ICD; this included the patient with VT on presentation who was implanted as secondary prevention. Four others had primary prevention devices.

### 2. Localisation of VT breakout

In 6 BrS (46%) and in 4 ARVC (36%) patients, VT was induced during the VT stimulation protocol. A consistent feature seen in all cases was progressive slowing of conduction, followed by formation of an arc of functional conduction block on the RVOT endocardium. Lines of functional block evolved on a beat-to-beat basis. In 5 cases, the VT then degenerated to VF. Figure 3 shows an example of a BrS patient with initiation of VT after development of an arc of functional block during a VT stimulation study (subject 6).

**Figure 3.**
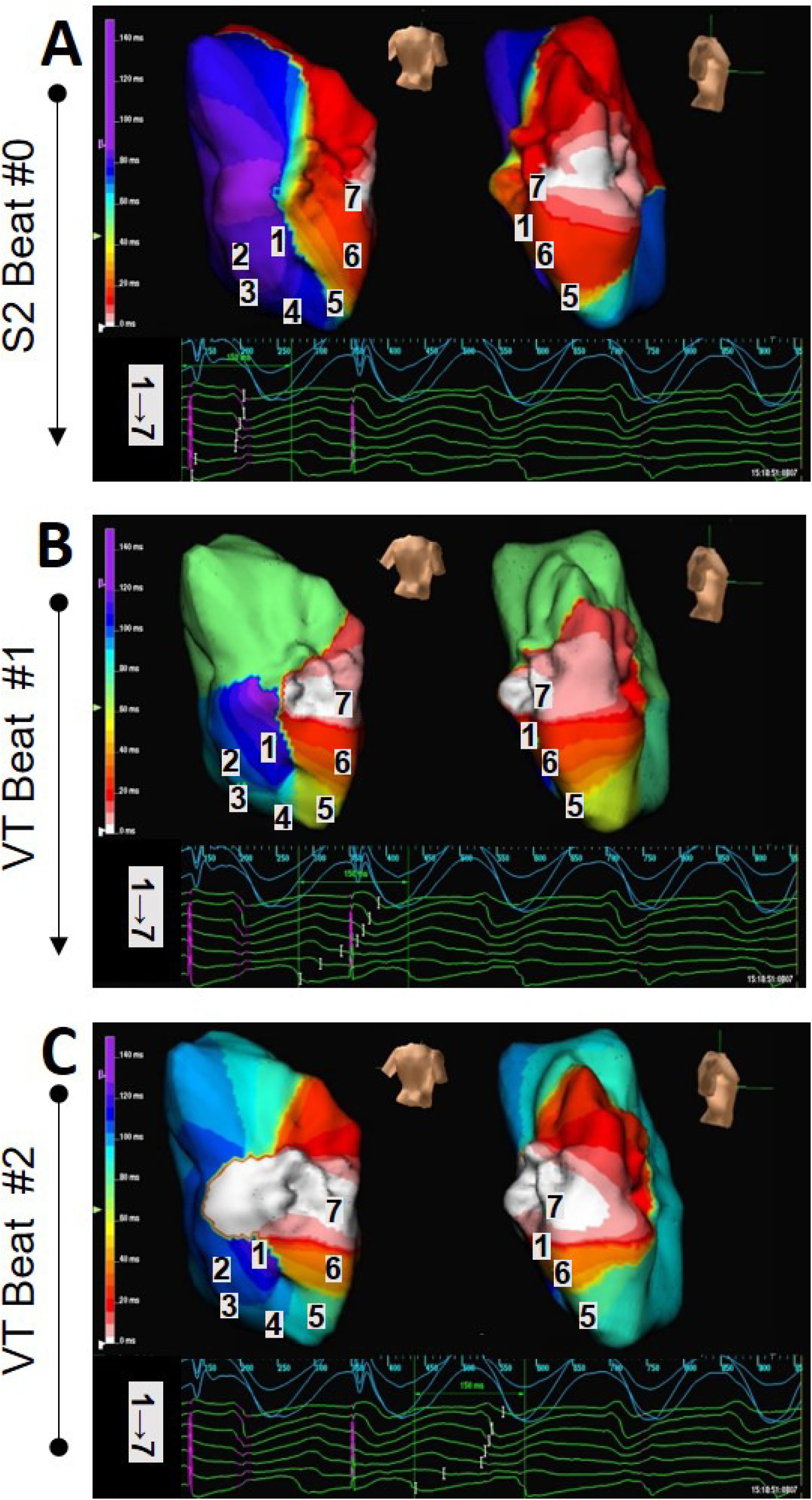
Geometry and isochronal RV maps created with the Ensite array in a BrS patient (subject 6), showing initiation of VT. The RV is show in posterior-anterior and right lateral views. The panels are immediately sequential in time. The time scale of the isochrones is shown on the left of each map. Sampled areas of reconstructed electrograms are shown in the lower panel, with the position of each electrogram marked on the isochronal maps. An S2 beat during a VT stimulation study results in a line of functional block (A). This results in the initiation of a re-entrant circuit around this persistent line of functional block on the posterior RV endocardium, with VT beat #1 shown in B and beat #2 shown in C.

Figure 4 shows representative spatial distributions of AT during VT and RVI calculated during the S1-S2 pacing protocol before VT. The shortest values of RVI, which represent sites of highest susceptibility to re-entry, co-localised with earliest activation point during VT in the ARVC (panel A) and BrS (panel B) patients where VT was initiated, but not in the patients with focal RVOT VT (panel C). The distance between region of lowest RVI and region of earliest activation during VT, *D_min_*, was significantly lower in BrS and ARVC than in focal VT (6.8 ± 6.7 ms vs 26.9 ± 13.3 ms, p=0.005) (Figure 5A). *D_min_* values were spread over a wide range of values in focal VT patients, with no clear association between region of lowest RVI and region of earliest activation during VT.

**Figure 4.**
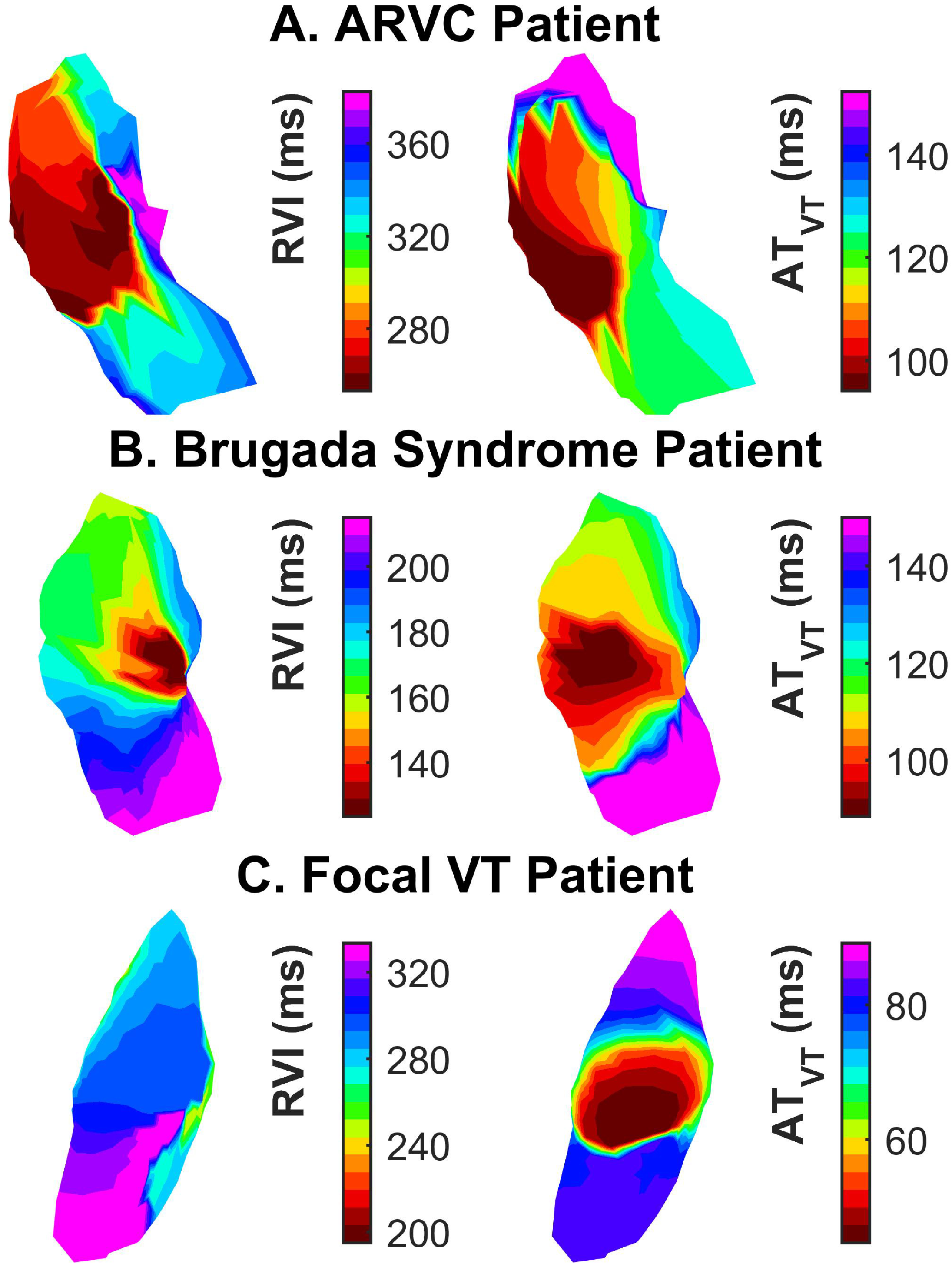
Spatial distributions of RVI measured before VT (right) and of AT during VT (left) from representative patients with ARVC (A), BrS (B) and focal VT (C), demonstrating co-localisation of regions of lowest RVI with VT breakout.

**Figure 5.**
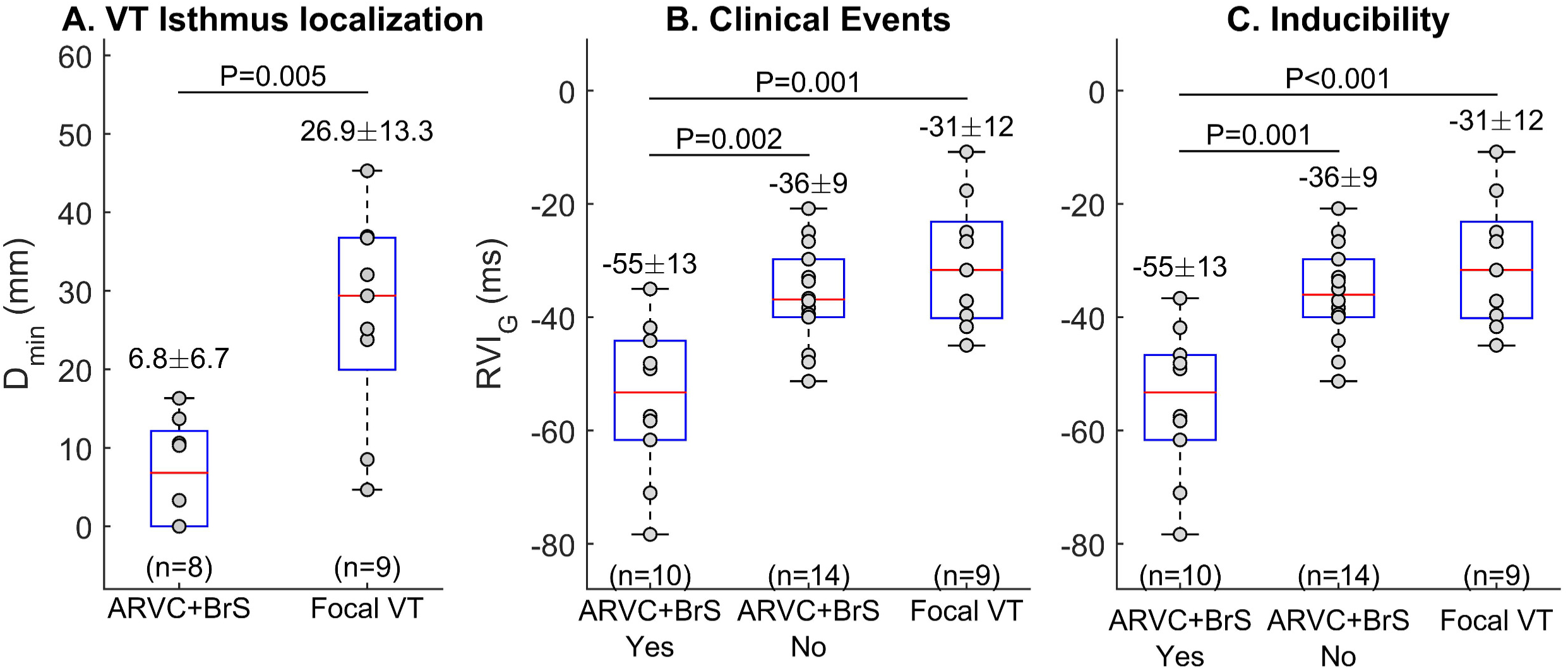
(A) Distance between region of lowest RVI and region of earliest activation during VT, *D_min_*, was significantly lower in BrS and ARVC than in focal VT. (B) BrS/ARVC patients with clinical VT events had lower global minimum RVI, RVI_G_, than those without VT or than patients with focal VT. (C) Patients with ARVC/BrS with inducible VT had significantly lower RVI_G_ than those with no VT or than patients with focal VT.

### 3. Low global RVI values predict likelihood of both clinical and inducible arrhythmias

Of the patients with ARVC/BrS, those with inducible VT had significantly lower RVI_G_ than those with no VT (-55.0 ± 13.0 ms vs -35.9 ± 8.6 ms, p= 0.001) (Figure 5). Patients with focal VT had the highest RVI_G_ (-30.6 ± 11.5 ms, p<0.001). The S_1_S_2_ coupling interval required for VT initiation was greater in those patients with clinical events compared to those without clinical events (298.6 ± 54.1 ms vs 213.3 ± 4.7, p=0.029). There was no significant difference in the routine clinical VERP estimation (217.8 ± 22.5 ms vs 214.0 ± 14.5, p=0.61) in inducible versus non-inducible patients.

Patients were followed up for 112 ± 19 months. Those with clinical VT events had lower RVI_G_ than those without VT (-54.5 ± 13.5 ms vs -36.2 ± 8.9 ms, p=0.002) or than those with focal VT (-30.6 ±11.5 ms, p=0.001) (Figure 5). Clinical events in our cohort included VF (2), haemodynamically unstable VT that terminated spontaneously (4) or was successfully treated by ICD with anti-tachycardia pacing (ATP) (3).

## Discussion

The results of this study show that RVI, a metric based on relative local ATs and RTs, can be employed in the analysis of re-entrant arrhythmias occurring in ARVC and BrS patients. The algorithm localizes regions of high susceptibility to conduction block and re-entry, with lowest RVI values identifying the breakout of re-entrant but not focal arrhythmias and it associates low global RVI with a propensity for clinically significant ventricular arrhythmias. This was identified rapidly utilising non-contact mapping.

The identification of optimal targets for catheter ablation of scar-related VT remains a significant challenge. Catheter ablation therapies usually target the exit (or entry) site of the slow-conducting isthmuses of viable myocardium that often transect dense scar (13, 14). However, accurate identification of the exit sites often requires VT induction, increasing the risk of the procedure as the VT may not be haemodynamically tolerated. The VT may also be difficult to induce or sustain, and may be mechanistically different to spontaneously occurring clinical VT. Voltage-mapping or detection of fractionated electrograms is frequently used to locate the isthmus region, but these techniques do not consider the repolarisation landscape around the scar and consequently find it difficult to identify the critical exit sites forming part of the re-entrant circuit, often leading to later recurrence. Furthermore, due to the often extensive nature of scar, large areas of ventricular myocardium are ablated using a “substrate based” approach leading to long procedures and risking increased myocardial damage. There is therefore a strong case for using a novel functional as opposed to anatomical substrate mapping technique incorporating both AT and RT information to predict sites of VT breakout without inducing VT. This could be further enhanced utilising a noncontact mapping approach to globally interrogate the ventricle. In our study, sites of lowest RVI did not correspond to the site of VT breakout in focal VT. This is in keeping with a mechanism of cAMP mediated triggered activity as opposed to localised re-entry (15).

Increasingly, ablation is being used as a treatment option of inherited arrhythmic conditions. The majority of patients presenting with VT secondary to BrS and ARVC tend to have complex scar morphologies with patchy fibrosis and a number of different conduction pathways/channels that may support re-entrant circuits with multiple potential exit sites (16–19, 2). However, previous work from our group has shown that both BrS and ARVC are distinguished by marked decremental conduction delays even in the absence of detectable fibrosis on imaging, indicating that functional electrophysiological changes are critical to the initiation of VT in these patients, especially early in the disease process (20–22). The proposed methodology can successfully identify exit sites in these conditions corresponding to the induced re-entrant VT circuits.

A further source of contention is risk stratification for the inherited arrhythmic conditions. SCD is often the first evidence of disease; therefore, identification of patients at highest risk is of utmost importance. Autopsy series and observational studies have reported a number of independent predictors of adverse outcomes in ARVC, in particular a prior history of sustained VT/VF and RV or LV dysfunction (23, 24); however these often occur late in the disease. Other potential risk factors with limited evidence include QRS fragmentation (25), fragmented electrograms (26), extent of T-wave inversion (27) and precordial QRS amplitude ratio (25). Recently AF, syncope, strenuous exercise after diagnosis, haemodynamically tolerated monomorphic VT and male sex have been reported as important risk factors for life threating arrhythmia (28). There is conflicting data regarding the role of an electrophysiology study in risk stratification of SCD in ARVC (21-23). Similarly, while the high-risk BrS patient is recognized through aborted SCD or unexplained syncope (29), risk stratification in asymptomatic patients is ill-defined, with inconsistent results from both non-invasive and invasive prognostic tests. These include spontaneous type 1 pattern, *SCN5A* mutation, syncope, family history of SCD, inducible arrhythmias, VERP < 200ms, QRS fragmentation, history of atrial fibrillation, lead I S-wave amplitude and duration (29–31).

It is logical that a metric which incorporates both repolarisation and conduction changes may accurately reflect the potential for arrhythmogenesis in both these conditions. The RVI algorithm was able to distinguish between patients with ARVC and BrS who had haemodynamically compromising clinical arrhythmias over nearly a decade of follow-up, as well as distinguishing re-entrant from focal focal VT. This index could thus potentially be applied to risk stratify ARVC/BrS patients to target ICD prophylaxis or indeed give an indication of early cardiomyopathic process in RVOT ectopic cases. Development of the analysis software for real-time signal processing could then allow establishment of this clinical tool for integration into existing clinical EP systems.

## Limitations

We paced from the RV apex; however, it is known that if the pacing site is changed, this may shift activation and repolarisation (11) and may in some cases shift exit sites for VT. Simulations in ischaemic cardiomyopathy have demonstrated that the RVI map may also change (7) and criteria for the selection of optimal pacing site should be identified in further studies. Our study population was relatively small, and to enable clinical translation, prospective testing in a larger cohort would be necessary to enable clinical translation and demonstrate its superiority over other common substrate mapping procedures. This is the subject of ongoing work. VT circuits were mapped in this study with non-contact activation maps; confirmation of the VT isthmus by entrainment was not possible as the VTs were haemodynamically unstable.

## Conclusions

RVI mapping has the potential to significantly advance both risk stratification and VT ablation therapy in ARVC and BrS, where inducibility of VT at EPS remains highly contentious and additional risk markers are needed, and may provide specific ablation targets for non-inducible VT without the need for full substrate ablation.

## Perspectives

Competency in Medical Knowledge: Unidirectional block and re-entry are the major determinants of VT initiation. VTs are often unstable and difficult to map sequentially due to multiple morphologies and haemodynamic compromise-employing a new metric (RVI) to predict sites of re-entry from local conduction and repolarisation parameters without inducing VT identifies the isthmus of VT and predicts probability of future events in ARVC and BrS.

Translational Outlook: Utilising conduction and repolarisation parameters without having to induce VT has important implications in identifying sites for focused ablation and risk stratification for ICD implantation. This approach needs to be evaluated in a prospective randomised study.

## References

1. Basso C, Bauce B, Corrado D, Thiene G. Pathophysiology of arrhythmogenic cardiomyopathy. Nat. Rev. Cardiol. 2011; 9: 223–233.

2. Nademanee K, Raju H, de Noronha SV, et al. Fibrosis, Connexin-43, and Conduction Abnormalities in the Brugada Syndrome. J. Am. Coll. Cardiol. 2015; 66: 1976–1986.

3. Papavassiliu T, Veltmann C, Doesch C, et al. Spontaneous type 1 electrocardiographic pattern is associated with cardiovascular magnetic resonance imaging changes in Brugada syndrome. Heart Rhythm 2010; 7: 1790–1796.

4. Opthof T, Janse MJ, Meijborg VMF, Cinca J, Rosen MR, Coronel R. Dispersion in ventricular repolarization in the human, canine and porcine heart. Prog. Biophys. Mol. Biol. 2016; 120: 222–235.

5. Coronel R, Wilms-Schopman FJG, Opthof T, Janse MJ. Dispersion of repolarization and arrhythmogenesis. Heart Rhythm Off. J. Heart Rhythm Soc. 2009; 6: 537–543.

6. Child N, Bishop MJ, Hanson B, et al. An activation-repolarization time metric to predict localized regions of high susceptibility to reentry. Heart Rhythm Off. J. Heart Rhythm Soc. 2015; 12: 1644–1653.

7. Hill YR, Child N, Hanson B, et al. Investigating a Novel Activation-Repolarisation Time Metric to Predict Localised Vulnerability to Reentry Using Computational Modelling. PloS One 2016; 11: e0149342.

8. Hindricks G, Kottkamp H. Simultaneous noncontact mapping of left atrium in patients with paroxysmal atrial fibrillation. Circulation 2001; 104: 297–303.

9. Dixit S, Lavi N, Robinson M, et al. Noncontact electroanatomic mapping to characterize typical atrial flutter: participation of right atrial posterior wall in the reentrant circuit. J. Cardiovasc. Electrophysiol. 2011; 22: 422–430.

10. Orini M, Taggart P, Srinivasan N, Hayward M, Lambiase PD. Interactions between Activation and Repolarization Restitution Properties in the Intact Human Heart: In-Vivo Whole-Heart Data and Mathematical Description. PloS One 2016; 11: e0161765.

11. Srinivasan NT, Orini M, Simon RB, et al. Ventricular stimulus site influences dynamic dispersion of repolarization in the intact human heart. Am. J. Physiol. Heart Circ. Physiol. 2016;311:H545–554.

12. Hanson B, Sutton P, Elameri N, et al. Interaction of activation-repolarization coupling and restitution properties in humans. Circ. Arrhythm. Electrophysiol. 2009; 2: 162–170.

13. Tung R, Boyle NG, Shivkumar K. Catheter ablation of ventricular tachycardia. Circulation 2010;122: e389–391.

14. Stevenson WG, Soejima K. Catheter Ablation for Ventricular Tachycardia. Circulation 2007; 115: 2750–2760.

15. Lerman BB, Stein K, Engelstein ED, et al. Mechanism of repetitive monomorphic ventricular tachycardia. Circulation 1995; 92: 421–429.

16. Nademanee K, Veerakul G, Chandanamattha P, et al. Prevention of ventricular fibrillation episodes in Brugada syndrome by catheter ablation over the anterior right ventricular outflow tract epicardium. Circulation 2011; 123: 1270–1279.

17. Zhang P, Tung R, Zhang Z, et al. Characterization of the epicardial substrate for catheter ablation of Brugada syndrome. Heart Rhythm Off. J. Heart Rhythm Soc. 2016.

18. Haqqani HM, Tschabrunn CM, Betensky BP, et al. Layered activation of epicardial scar in arrhythmogenic right ventricular dysplasia: possible substrate for confined epicardial circuits. Circ. Arrhythm. Electrophysiol. 2012; 5: 796–803.

19. Calvo D, Atienza F, Saiz J, et al. Ventricular Tachycardia and Early Fibrillation in Patients With Brugada Syndrome and Ischemic Cardiomyopathy Show Predictable Frequency-Phase Properties on the Precordial ECG Consistent With the Respective Arrhythmogenic Substrate. Circ. Arrhythm. Electrophysiol. 2015; 8: 1133–1143.

20. Lambiase PD, Ahmed AK, Ciaccio EJ, et al. High-density substrate mapping in Brugada syndrome: combined role of conduction and repolarization heterogeneities in arrhythmogenesis. Circulation 2009;120:106–117, 1–4.

21. Gomes J, Finlay M, Ahmed AK, et al. Electrophysiological abnormalities precede overt structural changes in arrhythmogenic right ventricular cardiomyopathy due to mutations in desmoplakin-A combined murine and human study. Eur. Heart J. 2012; 33: 1942–1953.

22. Finlay MC, Ahmed AK, Sugrue A, et al. Dynamic Conduction and Repolarisation Changes in Early Arrhythmogenic Right Ventricular Cardiomyopathy versus Benign Outflow Tract Ectopy Demonstrated by High Density Mapping & Paced Surface ECG Analysis. PLoS ONE 2014;9. Available at: http://www.ncbi.nlm.nih.gov/pmc/articles/PMC4094482/. Accessed October 21, 2016.

23. Link MS, Laidlaw D, Polonsky B, et al. Ventricular Arrhythmias in the North American Multidisciplinary Study of ARVC. J. Am. Coll. Cardiol. 2014; 64: 119–125.

24. Corrado D, Wichter T, Link MS, et al. Treatment of Arrhythmogenic Right Ventricular Cardiomyopathy/Dysplasia: An International Task Force Consensus Statement. Circulation 2015; 132: 441–453.

25. Saguner AM, Ganahl S, Baldinger SH, et al. Usefulness of Electrocardiographic Parameters for Risk Prediction in Arrhythmogenic Right Ventricular Dysplasia. Am. J. Cardiol. 2014; 113: 1728–1734.

26. Santangeli P, Dello Russo A, Pieroni M, et al. Fragmented and delayed electrograms within fibrofatty scar predict arrhythmic events in arrhythmogenic right ventricular cardiomyopathy: Results from a prospective risk stratification study. Heart Rhythm 2012; 9: 1200–1206.

27. Bhonsale A, James CA, Tichnell C, et al. Risk Stratification in Arrhythmogenic Right Ventricular Dysplasia/Cardiomyopathy-Associated Desmosomal Mutation Carriers. Circ. Arrhythm. Electrophysiol. 2013; 6: 569–578.

28. Mazzanti A, Ng K, Faragli A, et al. Arrhythmogenic Right Ventricular Cardiomyopathy: Clinical Course and Predictors of Arrhythmic Risk. J. Am. Coll. Cardiol. 2016; 68: 2540–2550.

29. Kawazoe H, Nakano Y, Ochi H, et al. Risk stratification of ventricular fibrillation in Brugada syndrome using noninvasive scoring methods. Heart Rhythm 2016; 13: 1947–1954.

30. Probst V, Veltmann C, Eckardt L, et al. Long-term prognosis of patients diagnosed with Brugada syndrome: Results from the FINGER Brugada Syndrome Registry. Circulation 2010; 121: 635–643.

31. Priori SG, Gasparini M, Napolitano C, et al. Risk stratification in Brugada syndrome: results of the PRELUDE (PRogrammed ELectrical stimUlation preDictive valuE) registry. J. Am. Coll. Cardiol. 2012; 59: 37–45.

